# Molecular mechanism for kinesin-1 direct membrane recognition

**DOI:** 10.1101/2021.01.20.427326

**Authors:** Zuriñe Antón, Johannes F. Weijman, Christopher Williams, Edmund R.R. Moody, Judith Mantell, Yan Y. Yip, Jessica A. Cross, Tom A. Williams, Roberto A. Steiner, Matthew Crump, Derek N. Woolfson, Mark P. Dodding

## Abstract

The cargo-binding capabilities of cytoskeletal motor proteins have expanded during evolution through both gene duplication and alternative splicing. For the light chains of the kinesin-1 family of microtubule motors, this has resulted in an array of carboxy-terminal domain sequences of unknown molecular function. Here, combining phylogenetic analyses with biophysical, biochemical and cell biology approaches we identify a highly conserved membrane-induced curvature-sensitive amphipathic helix within this region of a newly defined subset of long kinesin light chain paralogues and splice isoforms. This helix mediates the direct binding of kinesin-1 to lipid membranes. Membrane binding requires specific anionic phospholipids and is important for kinesin-1 dependent lysosome positioning, a canonical activity that until now has been attributed exclusively the recognition of organelle-associated cargo adaptor proteins. This leads us to propose a new protein-lipid coincidence detection framework for kinesin-1 mediated organelle transport.

## Introduction

While ancestral reconstructions suggest that early eukaryotes were already capable of kinesin-mediated microtubule transport, the kinesin repertoire has been elaborated throughout eukaryotic evolution through gene duplication and alternative splicing^1,2^. How this complexity subsequently confers broader functional capacity often remains unclear. This question is particularly pertinent for the kinesin-1 family that uses a remarkably versatile cargo recognition mechanism to enable the binding and transport of protein and ribonuclear protein complexes, viruses, and many different membrane-bound organelles (MBO)^3–5^. This mechanism comprises an interlinked and co-operative network of cargo binding and autoregulatory elements within the motor ATPase bearing kinesin heavy chains (KHCs) and kinesin light chains (KLCs)^6–8^.

The number of KLC genes varies between species; humans and mice have four paralogous KLCs (KLC1-4)^9–18^. KLC pairs combine with one of three KHC dimers (encoded by the KIF5A, B or C paralogues) to form an array of kinesin-1 heterotetramers with different cell and tissue expression profiles and cargo binding properties^19–23^. The physiological importance of kinesin-1 composition is underscored by a range of different neurological diseases that are reported to be contributed to or caused by dysregulation of particular KHCs and KLCs. These include loss of KLC4 function in hereditary spastic paraplegia, upregulation of KLC1 isoform E (KLC1E) in Alzheimer’s Disease (AD) and upregulation of KLC2 in spastic paraplegia, optic atrophy and neuropathy (SPOAN) syndrome^24–27^.

KLCs recognise MBO-associated cargo adaptor proteins using their highly conserved tetratricopeptide repeat domains (KLC^TPR^). Recent structural studies have revealed the molecular details of this motor-cargo interface that can engage both leucine-zipper and short-linear peptide motif (SLiM) features in adaptor proteins. These adaptors act as a regulatory tether between motor and cargo that couples cargo binding with motor activity and transport^7,28–31^. KLC amino acid sequences mainly diverge in their carboxy-terminal domains (KLC^CTD^: defined here as any protein sequences after the end of KLC^TPR^). These help to direct MBO-targeting^32–34^ and enable the phosphorylation-dependent regulation of kinesin-1 activity in axonal transport^35,36^. The molecular basis for these activities is not understood.

Here we describe the discovery and characterization of an evolutionarily conserved, membrane-induced, amphipathic helix within the CTDs of a newly defined class of long KLCs that mediates the direct phospholipid dependent binding of kinesin-1 to membranes and is important for the transport of lysosomes.

## Results

### *In silico* sequence, structural and phylogenetic analysis reveals an amphipathic helix within a newly defined class of long KLCs

A comparison of KLC^CTD^ primary sequences and their predicted secondary structures indicated the presence of a lone α helix within an otherwise unstructured domain of the long KLC1 splice isoforms (KLC1D and KLC1E, encoded by exon 16) as well as KLC2 and KLC4 but not the short KLC1 isoforms (KLC1 A, B, C, F, G) (Figure 1A,B). The predicted helix in KLC1D/E is also found in the KLC1 I, J, K, M, N and O variants that are not considered in the present study^15^. Further characterisation of this sequence suggested that the helix, if formed, would have a hydrophobic face and a net positively charged face (*i.e*., an amphipathic helix, AH) (Figure 1C). A shorter sequence that retains some similar hydrophobic and charged amino acids encoded by exon 14 (skipped in KLC1D/E) is found in KLC1B and KLC1C (Figure 1B). Further complexity arises in the case of KLC3; a structure-based alignment revealed that unlike the other paralogues KLC3^TPR^ terminates within TPR5, and so it lacks the final 6^th^ TPR (Supplementary Figure 1). However, the predicted AH is still present after the truncated KLC^TPR^ domain (Figure 1C, Supplementary Figure 1).

**Figure 1.**
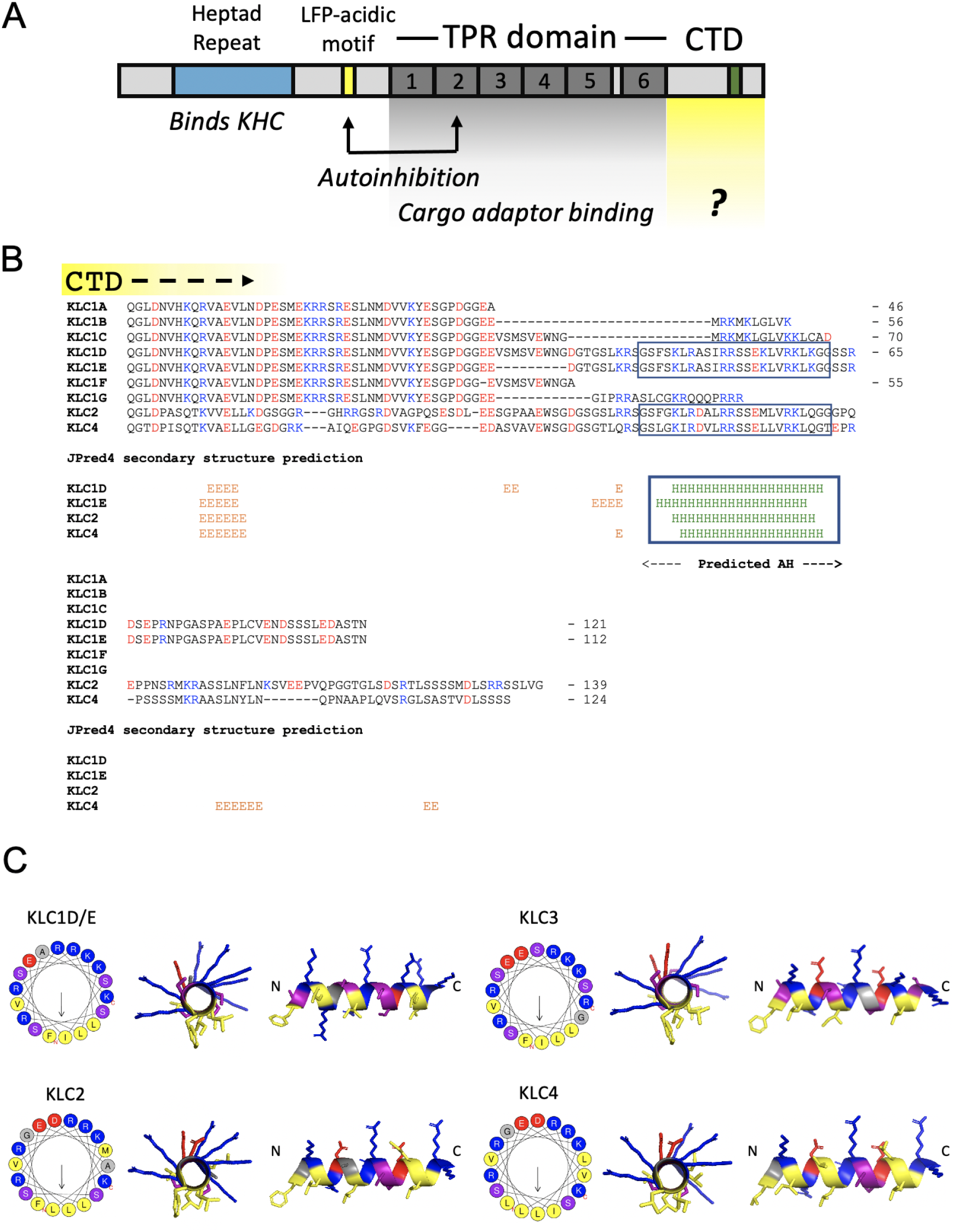
Long KLC paralogues and isoforms carry an amphipathic helix within their carboxy-terminal domain. (A) Domain architecture of the KLCs. (B) Manually edited PROMALS3D-based alignment and JPred4 secondary structure predictions for the CTDs of mouse KLCs 1-4. Lengths are indicated from a conserved post-KLC^TPR^ QG(L/T)D sequence. H-helix, E-extended. (C) Visualization of the AH in KLCs 1-4 using Heliquest and PyMOL.

We identified the same AH feature in vertebrate (chicken, frog, and zebrafish) KLC orthologues as well as the long sea urchin (*Stronglyocentrotus purpuratus*) KLC isoforms, SPKLC1-3, but not the short SPKLC4^14^ (Supplementary Figure 2). BLASTp searches of the NCBI non-redundant (nr) database indicated its presence in many other deuterostome KLCs (Supplementary Figure 3A). The AH was not apparent in the relatively short fly (*Drosophila melanogaster*,^18^) or worm (*Caenorhabditis elegans*,^17^) orthologues; this is despite the fact their KLCs have very well conserved KLC^TPR^ domains that enables clear definition of the KLC^CTD^ (Supplementary Figure 2). Further analysis using HH-pred failed to reveal any significant similarity (E-value < 1) to any protein in these species^37^. However, further database analysis and secondary structure prediction did identify closely related AH sequences in the KLC^CTD^ of other protostomes, including Brachiopoda and Annelida, as well as non-bilaterian animals, including Cnidaria, Ctenophora, and Placozoa (Supplementary Figure 3B).

A KLC phylogeny (Supplementary Figure 4) indicated that the 4 paralogous KLC protein coding genes of human and mouse arose from gene duplications on the gnathostome stem, with KLC1-4 being represented in the various classes that make up Eugnathostomata: Mammalia, Eureptillia, Amphibia, Actinopterygii and Chondrichthyes. The multiple KLC homologues of Petromyzon appear to be the result of independent, lineage-specific duplications, with the same mechanism resulting in *C. elegans* klc-1 and klc-2. The presence of clear KLC orthologues in animals and choanoflagellates, but not in more distant metazoans, indicates the KLC gene originated in their common choanozoan ancestor. Paralogues of KLC are present in some Fungi, Viridiplantae and members of the SAR clade of eukaryotes. As for *D. melanogaster* and *C. elegans*, the AH region was not detected in sequences of the wider Ecdysozoa. This can be explained by secondary loss because the AH motif is present in non-bilaterian holozoan sequences, including choanoflagellates. The onset of alternative splicing as a means of producing AH-containing and AH-lacking protein variants in mice and humans likely arose on the agnathan stem lineage because the deep-branching agnathan *Petromyzon* (sea lamprey) possesses isoforms displaying the short and long form, whereas more distantly related genomes do not.

Together, these data show that the KLC protein family can be newly divided into short forms that lack, and long forms that carry, an evolutionarily well conserved ≈ 20 amino acid predicted AH within their CTD. In general, amino acids on the hydrophobic face are highly conserved whereas the abundance, position and polarity of charges vary on the hydrophilic face but retain overall basicity. Serine residues occur frequently but their number and position varies (Figure 1C, Supplementary Figures 2 and 3).

### The AH peptide adopts a helical conformation in the presence of model membranes

A synthetic peptide encompassing this sequence from KLC1D/E was analysed in vitro using circular dichroism (CD) spectroscopy (Figure 2A). In aqueous solution the KLC1D/E peptide was unstructured. However, in the presence of the zwitterionic membrane mimetic dodecylphosphocholine (DPC) micelles the CD spectrum changed to one characteristic of a highly folded α-helix (Figure 2A). To test this further, a control peptide with two aliphatic residues replaced by proline (IL/PP KLC1D/E) was designed to prevent helical folding. This gave a CD spectrum expected for a random coil, both in the absence and presence of DPC micelles (Figure 2A). Interestingly, the helical folding of the parent KLC1D/E peptide was not induced with the non-ionic detergents n-Dodecyl β-D-maltoside (DDM), Octyl β-D-glucopyranoside (OG), or the zwitterionic detergent Lauryldimethylamine oxide (LDAO), indicating that specific electrostatic protein-lipid interactions may be important for AH folding (Supplementary Figure 5A).

**Figure 2.**
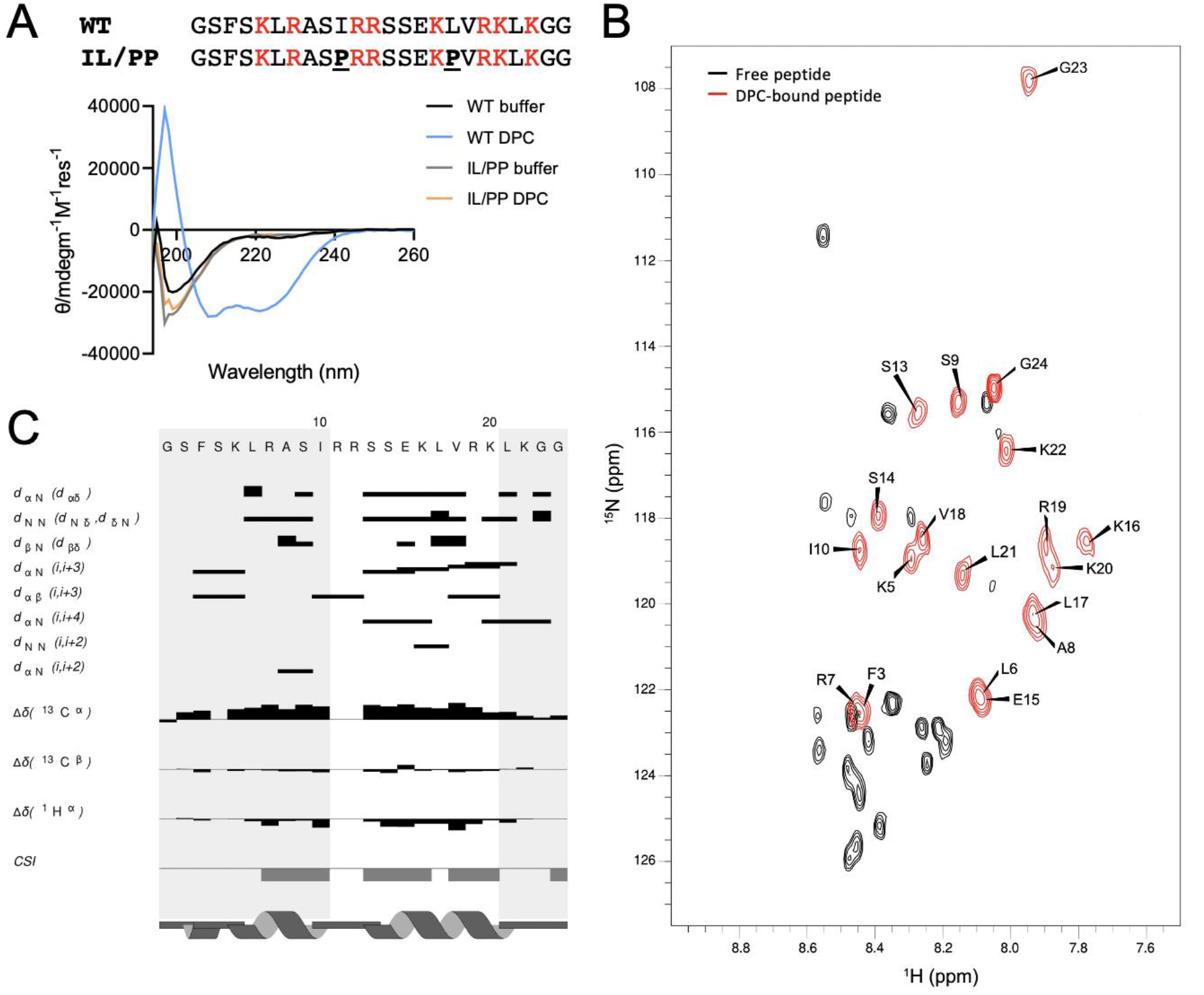
The AH peptide folds into a helix in the presence of model membranes. (A) Peptide sequences and their CD spectra in the absence or presence of 0.35% DPC. (B) Overlaid 15N HSQCs of the unbound (black) and DPC-bound (red with assignments) peptides. (C) NOE/CSI plot of the DPC-bound peptide sequence.

Next, we used nuclear magnetic resonance (NMR) to examine conformational changes in the peptide in more detail (Figure 2B,C and Supplementary Figure 5B,C). The 1D ^1^H NMR spectrum of the peptide in buffer was typical of a random coil with very little dispersion in either the amide or methyl regions (Supplementary Figure 5B). However, upon titration with perdeuterated DPC, the peptide chemical shifts became sharper and more dispersed, saturating at around 0.7% DPC. This was particularly evident in the amide region of spectrum indicative of folding in the presence of DPC (Supplementary Figure 5B). This was confirmed by collecting ^1^H-^15^N HSQC datasets at natural ^15^N abundance (Figure 2B). In the presence of deuterated DPC the ^1^H-^1^H NOESY and TOCSY spectra of the peptide gave sufficiently dispersed H_N_ and H_α_ resonances to allow a NOE-based sequential assignment of the spectra (Figure 2B). Backbone NH resonances could be assigned for ≈ 80% of the peptide with missing assignments at the N-terminus and in the central R11/R12 region (Figure 2B). Analysis of the 2D NOESY spectra identified a number of d_NN_(*i,i+1*) and d_αN_(*i,i+1*) throughout the peptide backbone along with several d_αN_(*i,i+3*) and d_αN_(*i,i+4*) NOE correlations. These are diagnostic of α-helical conformation (Figure 2C). In particular, a number d_αN_(*i,i+3*) and d_αN_(*i,i+4*) NOEs between residues S13 and K20 indicated clearly that this region is an α helix. Although there were distinct d_NN_(*i,i+1*), d_αN_(*i,i+1*) and d_αN_(*i,i+3*) NOEs in the N-terminus of the peptide, this region was less well defined due to greater overlap in the N_H_-H_α_ region of the 2D NOESY spectra. In addition to the NOE analysis the ^13^C_α_/C_β_ and H_α_ backbone chemical shifts for the assigned residues were extracted using a natural abundance ^1^H-^13^C HSQC. Subsequent chemical shift index (CSI) analysis of the chemical shifts for the DPC-bound peptide were again consistent with an α helix for two stretches of residues, K5 - I10 and S13 - K20, as observed in the NOESY spectra (Figure 2C).

### The AH mediates KLC^CTD^ binding to lipid membranes

To evaluate the capacity of KLC^CTD^ to bind membranes, we assembled liposomes [small (SUV) and large (LUV) unilamellar vesicles] from Folch Fraction I brain lipids and analysed the capacity of KLC^CTDs^ (Supplementary Figure 6A) to associate with them using co-sedimentation assays. We focussed on the KLC1 paralogue for these liposome binding assays because of the availability natural isoform controls. Purified recombinant KLC1E^CTD^ that contains the predicted AH (long) but not KLC1B^CTD^ (short) bound to LUVs in a liposome-concentration dependent manner (Figure 3A,B). Binding of KLC1E^CTD^ to SUVs was significantly greater than to LUVs indicating that the interaction is sensitive to membrane curvature (Figure 3B). In a separate experiment, as expected, KLC1A^CTD^ also failed to associate with membranes (Supplementary Figure 6B). A single helix-disrupting proline mutation (I560P in full length protein; I/P mutant) in the hydrophobic face of KLC1E^CTD^ AH was sufficient to eliminate any detectable membrane interaction (Figure 3C,D). Furthermore, amine cross-linking by BS3, in the absence or presence of LUVs and SUVs (Figure 3E) resulted in the AH- and liposome-dependent formation of KLC1E^CTD^ oligomers with electrophoretic mobility corresponding to dimers, trimers and tetramers as well as higher-order assemblies. Membranes were also remodelled because analysis of (non-crosslinked) samples by cryo-electron microscopy revealed extensive AH-dependent membrane bending and vesicle clustering as well as the accumulation of small vesicles (Figure 3F). We concluded that KLC1E^CTD^ binds directly to membranes and this interaction requires the AH.

**Figure 3.**
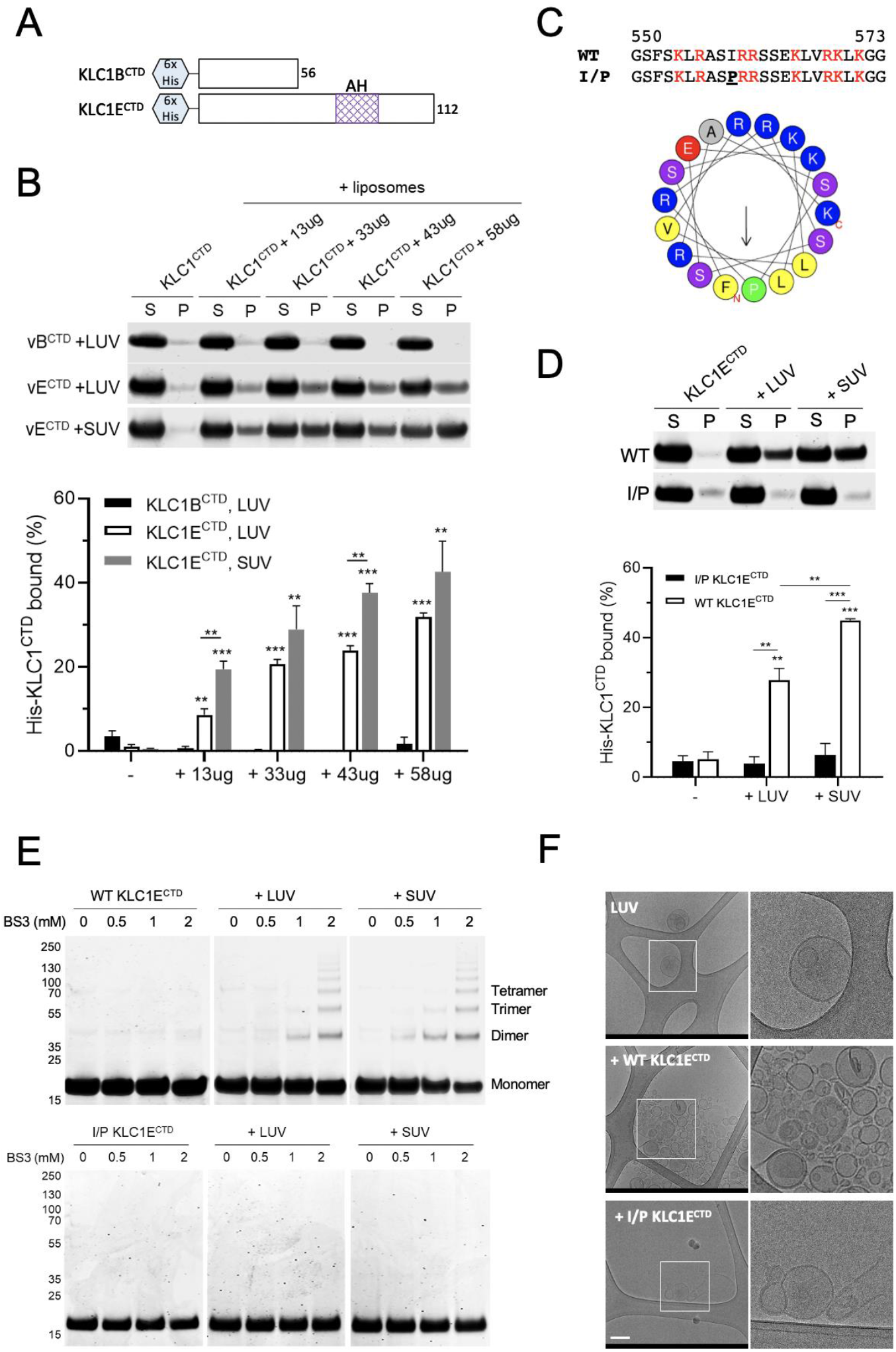
The AH is required for KLC1E^CTD^ binding to liposomes. (A) Representation of the His-tagged KLC1B/E^CTD^ constructs. (B) Co-sedimentation analysis of His-KLC1B/E^CTD^ with Folch fraction I LUV or SUV. Samples were analysed by SDS-PAGE/Coomassie (top) and quantified by densitometry (bottom). Means ± SEM of at least 3 experiments; **p<0.01; ***p<0.001 compared to the sample without liposomes or as indicated on the graph. (C) Sequence comparison and visualization of the AH in the I560P KLC1E^CTD^ mutant. (D) Co-sedimentation analysis of WT or I/P His-KLC1E^CTD^ with Folch fraction I LUV or SUV. Means ± SEM of at least 3 experiments; **p<0.01; ***p<0.001 compared to the sample without liposomes or as indicated in the graph. (E) BS3 crosslinking assay showing oligomers of WT but not I/P mutant His-KLC1E^CTD^ upon incubation with Folch fraction I LUV or SUV. (F) Cryo-EM observation of Folch fraction I LUV in the absence or presence of WT or I/P mutant KLC1E^CTD^. Scale Bar: 200 nm.

### Long KLC^CTD^ membrane binding is dependent on specific anionic phospholipids

Liposomes of defined composition were prepared from pure lipids. No binding was observed with either WT or the helix-disrupting mutant KLC1E^CTD^ upon incubation with liposomes consisting of PC, PC:PE (7:3) or PC:PE:PS (5:3:2) (Figure 4A) (where PC is phosphatidylcholine; PE, phosphatidylethanolamine; and PS, phosphatidylserine), suggesting that an additional component is required. We considered phosphatidic acid (PA) to be a good candidate because KLC1 (along with KIF5B) has been identified recently in a proteomic screen for PA-binding proteins^38^. Indeed, WT KLC1E^CTD^, but not the helix disrupting mutant bound to PC:PE:PA (5:3:2) liposomes although to a slightly lesser extent than to the Folch fraction liposomes (Figure 4A). Moreover, KLC1B^CTD^ showed no detectable binding to PA containing liposomes (Supplementary Figure 6C).

**Figure 4.**
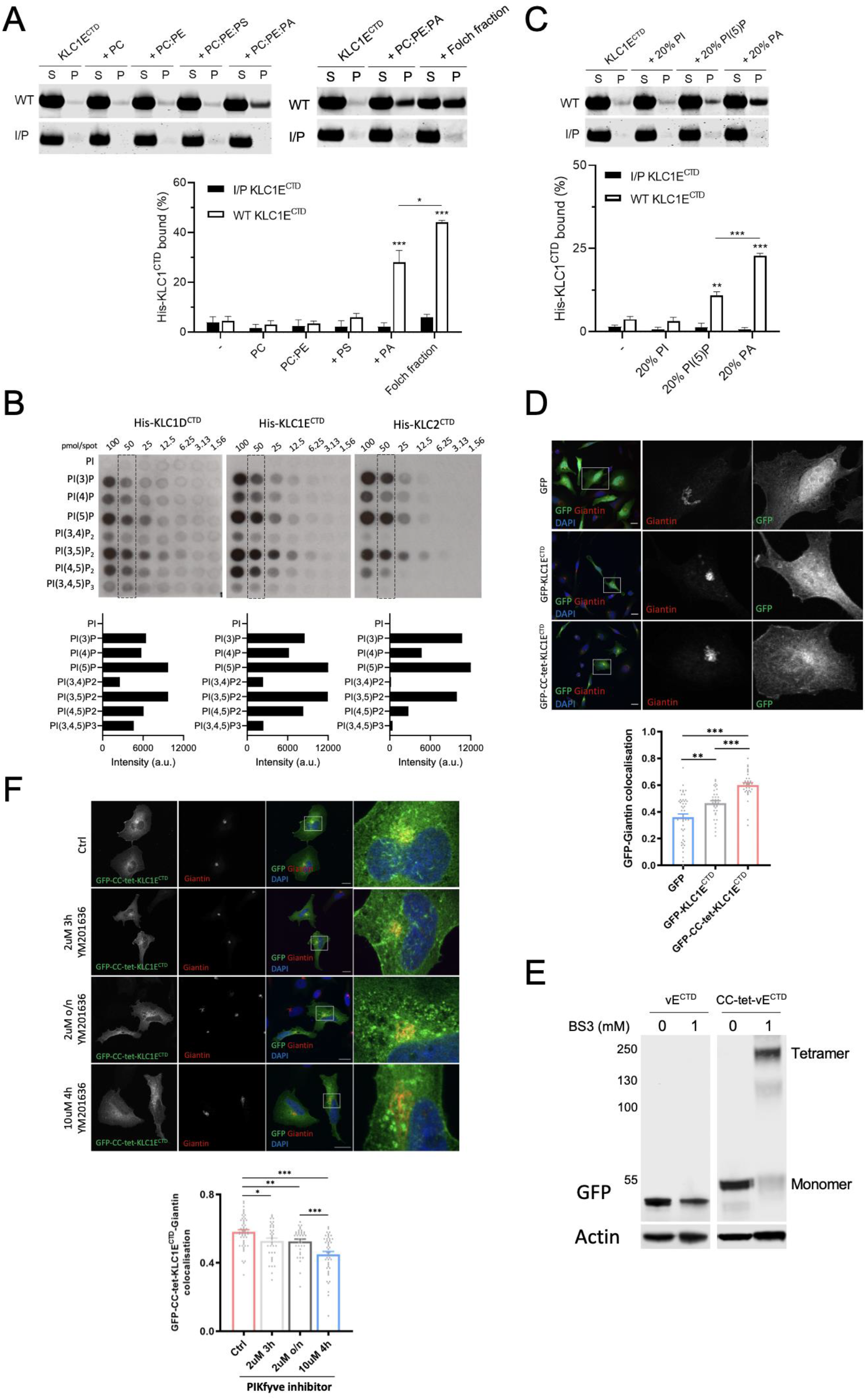
Long KLC^CTDs^ bind to specific anionic phospholipids. (A) Co-sedimentation analysis of WT or I/P His-KLC1E^CTD^ incubated with PC, PC:PE (7:3), PC:PE:PS (5:3:2), PC:PE:PA (5:3:2) or Folch fraction I LUV. (B) Binding of the indicated long KLC^CTDs^ to phosphoinositides in a protein lipid overlay assay. Quantitation is provided at the 50 pmol spot. (C) Co-sedimentation analysis of WT or I/P His-KLC1E^CTD^ incubated with PC:PE:PI, PC:PE:PI(5)P or PC:PE:PA (5:3:2) LUV. Means ± SEM of at least 3 experiments.; *p<0.05; **p<0.01; ***p<0.001 compared to the sample without liposomes or as indicated in the graph. (D) Immunofluorescence analysis (top) showing localization of GFP, GFP-KLC1E^CTD^, and GFP-CC-tet-KLC1E^CTD^. Golgi is stained with anti-giantin antibody. Representative Pearson’s correlation from one experiment and ≈ 40 cells (bottom). (E) BS3 crosslinking assay showing monomeric or tetrameric expression of KLC1E^CTD^ in lysates of HeLa cells transfected with GFP-KLC1E^CTD^ or GFP-CC-tet-KLC1E^CTD^. (F) GFP-CC-tet-KLC1E^CTD^-giantin co-localisation in cells treated with the PIKfyve inhibitor YM201636 as indicated (top). Representative Pearson’s correlation from one experiment and ≈ 40 cells (bottom). Mean ± SEM; *p<0.05; **p<0.01; ***p<0.001. Scale Bars: 20 μm.

Together, these data above show that KLC1E^CTD^ binds to PA in an AH dependent manner. However, as PA only comprises a small component of Folch fraction I liposomes, and even at 20% as used in our pure lipid assays was insufficient to match the degree of binding, we considered the possibility that KLC1E^CTD^ has a broader capacity for acidic phospholipid recognition. Consistent with this, long KLC^CTDs^ (KLC1D, KLC1E and KLC2) bound to phosphoinositides on membrane strips with a strong preference for PI(5)P and PI(3,5)P2, moderate association with PI(3)P, PI(4)P and PI(4,5)P2, very little binding to PI(3,4)P2 and PI(3,4,5)P3, and no detectable interaction with PI (Figure 4B). The short variants lacking the AH (KLC1A/B/C^CTD^) did not interact with any phosphoinositide (Supplementary Figure 6D). As these strips present the lipid outside of the context of a bilayer, we performed liposome binding assays with the top binding candidate - PI(5)P (Figure 4C). Consistent with the PIP array data, KLC1E^CTD^ was also capable of interacting with PI(5)P, but not PI, although to a lesser extent than with the same amount of PA (Figure 4C). Again, the helix-disrupting mutant showed no binding (Figure 4C). We concluded that the AH is essential for KLC1^CTD^ mediated membrane interaction, it shows some requirement and selectivity for specific anionic phospholipids, and that those properties extend to the closely related sequences in KLC2^CTD^.

To explore these interactions in cells, we compared the subcellular localisation of GFP and GFP-KLC1E^CTD^ in HeLa cells (Figure 4D). Whereas GFP was diffusely distributed throughout the cytosol and nucleus, GFP-KLC1E^CTD^ signal was slightly enriched in the perinuclear region. It partially co-localised with antibody staining for the Golgi marker giantin (Figure 4D), consistent with the previously reported Golgi localisation for the KLC1D/E isoforms^32,34^. We anticipated that forced oligomerisation of GFP-KLC1E^CTD^ might increase avidity and so enhance target membrane association. To test this, we introduced the sequence for a *de novo* designed coiled-coil tetramer between GFP and KLC1E^CTD^ (GFP-CC-tet-KLC1E^CTD^)^39^. The mobility of the fusion proteins from HeLa cell lysates was monitored by SDS-PAGE and western blotting (Figure 4E). Confirming the functionality of the coiled-coil sequence, a band shift to the expected molecular weight of an SDS-resistant tetramer was only observed with the GFP-CC-tet-KLC1E^CTD^ after BS3 treatment (Figure 4E). As predicted, the Golgi membrane association became more pronounced with GFP-CC-tet-KLC1E^CTD^, as determined by giantin co-localisation analysis of the GFP signal (Figure 4D); although there was clearly some recruitment to other compartments suggesting that KLC1E^CTD^-organelle recruitment is not highly specific. However, since the KLC^CTDs^ appeared to have limited preference for two 5’ phosphorylated phosphoinositides in vitro (PI(5)P and PI(3,5)P2), we used the PIKfyve kinase inhibitor YM201636 to reduce 5’ phosphorylation. This resulted in a time and inhibitor concentration dependent reduction in co-localisation of GFP-CC-tet-KLC1E^CTD^ with giantin and led to the redistribution of GFP-CC-tet-KLC1E^CTD^ to more dispersed puncta (Figure 4F). Thus, consistent with the membrane binding properties observed in vitro using recombinant protein (Figure 4A-C), KLC1E^CTD^ localisation is sensitive to membrane phosphoinositide composition in cells.

### Long KLC^CTD^-dependent membrane association of kinesin-1 is phosphoinositide sensitive and is important for lysosome positioning

To determine how membrane binding by isolated KLC^CTD^ extends to the complete tetrameric holoenzyme, we focussed on endogenously expressed kinesin-1 in biochemical assays, first seeking to establish the KLC1 and KLC2 expression profile in HeLa cells. These were transfected with plasmids to express full-length HA-tagged KLC1A, KLC1E and KLC2 and compared to non-transfected cells by SDS-PAGE and western blot analysis. An anti-HA antibody detected the over-expressed proteins, and consistent with their molecular weights, HA-KLC1E and HA-KLC2 migrated more slowly than HA-KLC1A (Figure 5A). Anti-KLC1 antibodies detected both overexpressed isoforms of KLC1 with similar efficiency, but not KLC2, and also a slightly faster migrating band in all samples that was confirmed to be an endogenous KLC1 isoform by siRNA depletion of KLC1 (Figure 5A). Similarly, anti-KLC2 antibodies specifically detected overexpressed HA-KLC2 but not HA-KLC1 and a slightly faster migrating band that was confirmed as endogenous KLC2 by siRNA knockdown (Figure 5A). Together these data indicate that most KLC1 in HeLa cells is a short isoform. In addition, the approximately equal expression of the HA-tagged proteins and difference in band intensity between endogenous and over-expressed proteins for the KLC1 and KLC2 specific antibodies led us to conclude that HeLa cells express similar amounts of KLC1 and KLC2 (roughly 40:60).

**Figure 5.**
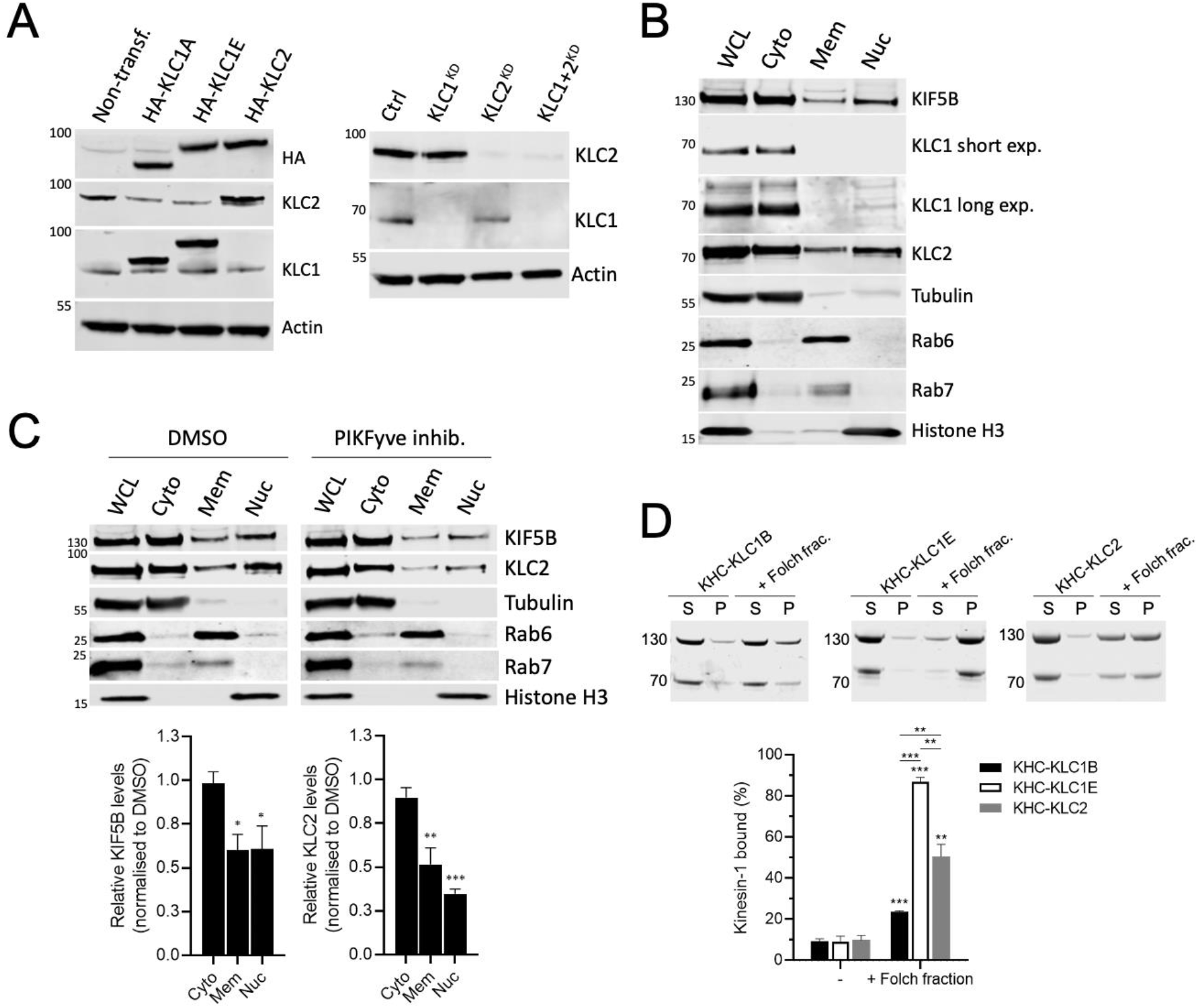
Membrane association of kinesin-1 is mediated by KLC^CTD^ and is phosphoinositide sensitive. (A) KLC1/2 expression in HeLa cells transfected with HA-KLC1A/1E/2 or non-transfected cells (left) or after siRNA knockdown of KLC1/2 (right) was analysed by western blot. (B) Fractionation and western blot analysis of endogenous KIF5B and KLC1/2. (C) Fractionation analysis of cells treated with DMSO or PIKfyve inhibitor YM201636 (10 μM, 4 h). Means ± SEM of at least 3 experiments; *p<0.05; **p<0.01; ***p<0.001 compared to the sample treated with DMSO. (D) Co-sedimentation assay and analysis by SDS-PAGE/Commassie staining using purified KHC-KLC1B/1E/2 and Folch fraction I LUV. Means ± SEM of at least 3 experiments; **p<0.01; ***p<0.001 compared to the sample without liposomes or as indicated in the graph.

To compare the localisation of the endogenous KLC paralogues, HeLa cell extracts were fractionated. As well as in the cytosolic fraction (containing β*-*tubulin), KLC2 was found in the membrane (containing Rab6 and Rab7) and nuclear (containing Histone H3) fractions (Figure 5B). In contrast, the shorter KLC1 was almost exclusively found in the cytosolic fraction (very weak bands in nuclear fraction on long exposure) and not at all in the membrane fraction, consistent with a specific role for the long KLCs membrane association. The major KHC paralogue in HeLa, KIF5B, was found in all three fractions (Figure 5B). Confirming a role for phosphoinositides in directing membrane association, treatment of cells with PIKfyve inhibitor reduced the amount of KLC2 and KIF5B in the membrane and nuclear fractions (Figure 5C).

To ask whether KLC^CTD^ is necessary and sufficient for direct membrane targeting of kinesin-1, KHC-KLC1B, KHC-KLC1E and KHC-KLC2 complexes were isolated from 293T cells by HaloTag purification and size-exclusion chromatography^8^. Consistent with a direct role in membrane binding of the holoenzyme, tetramers containing the long KLC1E or KLC2 sequences bound significantly better to Folch Fraction I liposomes in co-sedimentation assays compared to the short KLC1B isoform (Figure 5D).

PIKfyve inhibition modifies lysosome motility^40,41^ and lysosome distribution in HeLa is controlled by kinesin-1 through a well-defined cargo adaptor axis: BORC-Arl8-SKIP-KLC2-KIF5B, where SKIP acts as the cargo adaptor protein, binding KLC^TPR^ and KHC using the W-acidic class of SLiM ^42^. Indeed, siRNA knockdown of KLC2 caused reduction in the proportion of peripheral lysosomes compared to control cells (Figure 6A,B). A typical wildtype distribution profile characterised by many lysosomes in the cytoplasm outside the perinuclear region and accumulations at the cell periphery could be restored by expression of wildtype KLC2 but not KLC2 with a helix-disrupting mutation (L550P, equivalent to I560P in KLC1E). This confirms a role for the AH-membrane binding in kinesin-1 mediated lysosome positioning (Figure 6C,D). Similarly, expression of the long KLC1E but not the short KLC1A isoform restored cytoplasmic and peripheral lysosome accumulations. This indicates that the long AH containing paralogues have some functional redundancy in lysosome transport, consistent with their shared ancestry (Figure 6C,D).

**Figure 6.**
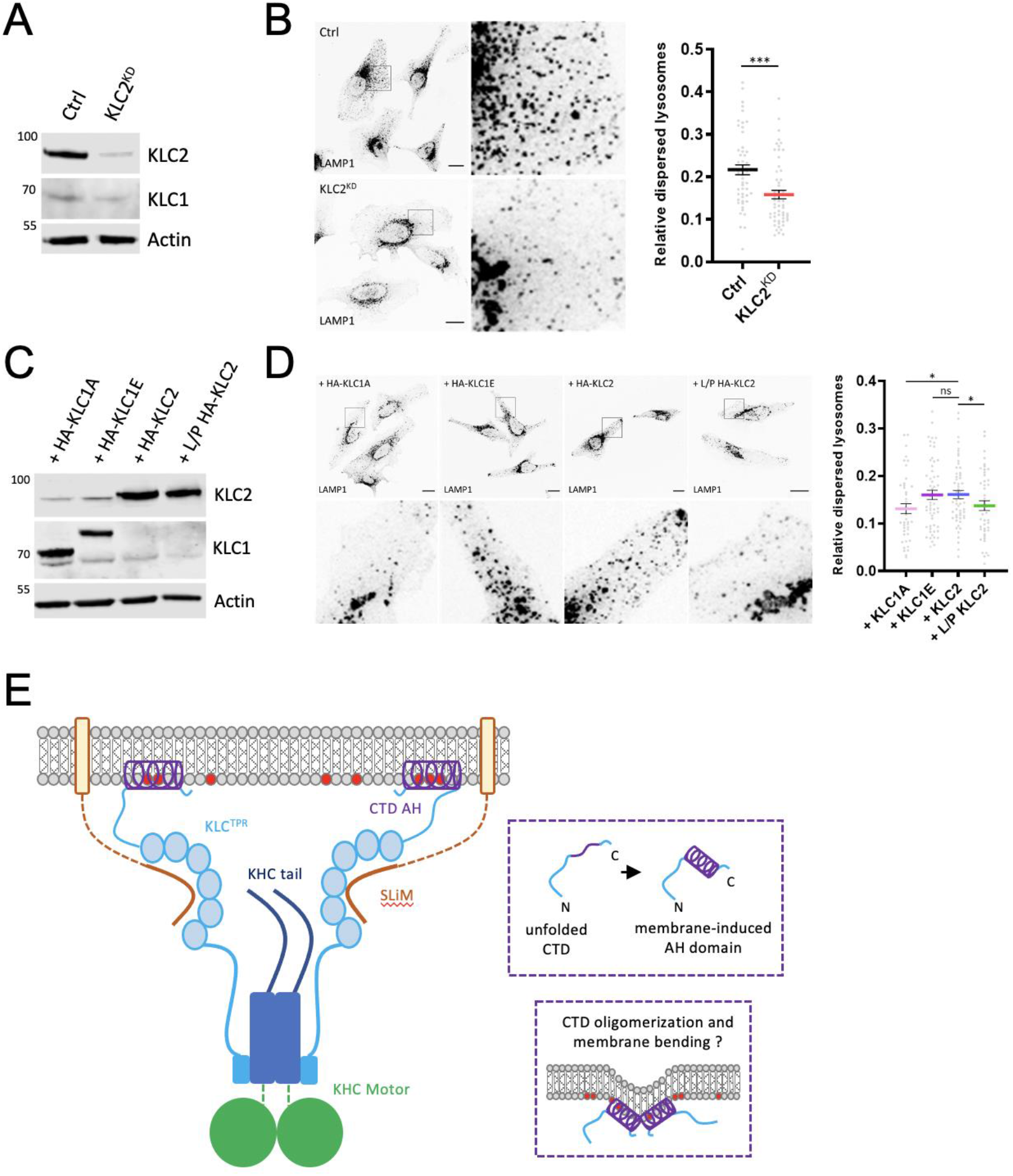
Long KLCs are required for kinesin-1 mediated lysosome positioning. (A) Western blot analysis of KLC2 expression in HeLa cells after siRNA knockdown of KLC2. (B) Lysosome distribution in control or KLC2 knockdown cells. Lysosome dispersion was determined by measuring perinuclear and whole cell LAMP1 signal fluorescence. Means ± SEM of at least 3 experiments; ***p<0.001. (C) KLC1/2 expression in KLC2 knockdown HeLa cells transfected with HA-KLC1A/1E/2/L550P 2 was analysed by western blot. (D) Lysosome distribution in KLC2 knockdown cells transfected with HA-KLC1A/1E/2/L550P 2. Means ± SEM with samples pooled from 3 independent experiments; *p<0.05. (E) Model showing representation of kinesin-1 cargo membrane association by the kinesin light chains. The membrane-induced helical folding and membrane remodeling capacity of the KLC^CTD^ helix are also represented.

## Discussion

The mechanistic basis for the critical role that the KLC^CTDs^ play in organelle transport has proved elusive. This is intimately linked with the question of why, particularly in vertebrates, the evolution of intracellular transport has favoured gene duplication and alternative splicing events that have resulted in so many KLCs that differ mainly in their CTD. The identification here of a membrane-induced AH within a newly defined subset of long KLC^CTDs^ provides an explanation, showing that the role of these sequences is to confer a direct membrane binding capacity on the motor complex. To the best of our knowledge, this is the first report of a membrane binding/induced AH being used directly by a microtubule motor protein. Importantly, the potential to attribute a relative increase in KLC1E or KLC2 expression to modification of the direct membrane properties of the kinesin-1 pool offers a pathway to understanding their reported roles in AD and SPOAN syndrome respectively^24,27^.

Given the well-established role of cargo adaptor proteins in motor recruitment and activation^6,7^, including at the lysosome, we favour a coat-like coincidence detection framework to explain our observations (Figure 6E). In this model, cargo adaptor proteins bind to KLC^TPR^ and membranes to KLC^CTD^. It will be important to sequence these protein and lipid interactions correctly. Previous work has indicated that the KLC^TPR^-adaptor binding gates access to cargo binding features of the KHC tail^43^. It seems reasonable to suggest that the KLC^CTD^-membrane interaction may function higher in this hierarchy to locally concentrate kinesin-1 at sites where it is available to engage cargo-adaptor proteins. It will be important to ask whether this involves curvature sensing to select target vesicles and if this is associated with membrane remodelling capacity. The nature of the final transport competent assembly will depend on the adaptor system because kinesin-1-cargo adaptors can be integral or peripheral membrane proteins^7^.

It is likely that phosphorylation will emerge as a key membrane binding regulator; CKII and GSK3β have been shown to phosphorylate KLC2^CTD^ to negatively regulate axonal transport and a serine residue within AH itself (S542 in mouse) as well as another at S579 in KLC2^CTD^ are phosphorylated by AMPK to inhibit axonal PI3K trafficking^35,36^. We posit that sequence differences between the long KLC^CTDs^ could offer the kinesin-1 family different sensitivity to particular kinases and phosphatases. A report of binding of 14-3-3 proteins to phosphorylated KLC2^CTD^ suggests that an AH/CTD-sequestration hypothesis is also worth considering as another level of regulation^44^.

What of the short KLCs? Our phylogenetic analysis suggests that membrane binding by the AH is an ancestral animal KLC feature. Counterintuitively, it appears that the loss of the AH rather than its acquisition may be associated with diversification of function. Previous reports have implicated KLC1B and KLC1C isoforms in rough endoplasmic reticulum and mitochondrial targeting despite lacking the exon 16 sequence encoded by the AH^15,32,33^. However, both isoforms share some sequences with limited homology to the second half of the AH (encoded by exon 14). Although helical properties of this 10-amino acid sequence were not confidently predicted by our *in silico* analysis, if folded, the resulting structure would be amphipathic. Whether it has a membrane/lipid binding capacity that was not detected in our assays is clearly worth further investigation. More generally, given their *de facto* positioning at the motor-organelle interface, there is a good case for a wider exploration of the membrane binding properties of the long and short KLC^CTDs^ throughout the wider KLC family. Specifically, it will be important to consider why the AH sequence was lost (relatively recently in evolutionary terms) in Ecdysozoa, including *D. melanogaster* and *C. elegans* and if KLC intrinsic or extrinsic compensatory mechanisms have emerged. It will also be interesting to ask whether some of the short KLC1 isoforms might be better adapted for the transport of non-membranous cargoes such as ribonuclear protein complexes or viral capsids^5,45^.

In conclusion, the identification of a membrane binding AH in the kinesin-1 cargo binding mechanism should lead to a step change in the view of how the family is classified, and cargo are recognised by this central player in intracellular transport. It points to a new direct role for membrane biology in KLC-dependent vesicle and organelle recognition that may be as important as cargo adaptor protein interactions themselves.

## Acknowledgments

This work was funded by the Biotechnology and Biosciences Research Council (BB/S000917/1 and BB/S000828/1). M.P.D. is a Lister Institute of Preventative Medicine Prize Fellow. We are grateful for the assistance and access to equipment at the Wolfson Bioimaging Facility and NMR Facility at the University of Bristol and BrisSynBio, a BBSRC/EPSRC-funded Synthetic Biology Research Centre (BB/L01386X/1). J.A.C is supported by the EPSRC Bristol Centre for Doctoral Training in Chemical Synthesis. E.R.R.M. is supported by a Royal Society Enhancement Award to T.A.W.. Y.Y.Y. was also supported by the Biotechnology and Biosciences Research Council (BB/L006774/1).

## Supplementary Materials

## Materials and Methods

### Reagents and antibodies

1,2-dioleoyl-sn-glycero-3-phosphoethanolamine (DOPE), bovine liver phosphatidylinositol (PI), brain phosphatidylserine (PS) and egg phosphatidic acid (PA) were purchased from Avanti Polar Lipids. Phosphatidylinositol 5-phosphate diC16 (PI(5)P) was obtained from Echelon Biosciences. Folch fraction I and egg phosphatidylcholine (PC) were purchased from Sigma-Aldrich. YM201636 PIKfyve inhibitor was purchased from Cayman Chemical. Cell fractionation kit was obtained from Cell Signalling. Non-targeting control siRNA was obtained from Horizon Discovery and human KLC1/2 siRNAs were from Santa Cruz Biotechnology. The following primary antibodies were used: HRP-conjugated anti-His6 (71841, Novagen), anti-GFP for immunoblotting (3E1, Roche), anti-actin (AC-74, Sigma-Aldrich), anti-beta-tubulin (AA2, Sigma-Aldrich), anti-histone H3 (Abcam), anti-giantin (PRB-114C, Covance), anti-HA (HA-7, Sigma-Aldrich), anti-KLC1 ([EPR12441(B)], Abcam), anti-KLC2 (Abcam), anti-KIF5B (Abcam), anti-LAMP1 (D2D11, Cell Signalling), anti-Rab7 (D95F2, Cell Signalling) and anti-Rab6 (D37C7, Cell Signalling). Alexa 568-, and 633-conjugated anti-mouse or anti-rabbit secondary antibodies were from Thermo Scientific.

### Peptides and plasmids

KLC1D/E synthetic peptides used for CD or NMR measurements were purchased from Genscript (>98% purity). Sequences were as follows: WT KLC1D/E, GSFSKLRASIRRSSEKLVRKLKGG; IL/PP KLC1D/E, GSFSKLRASPRRSSEKPVRKLKGG. Peptides were dissolved in water and stored at −20°C until use.

KLC1 isoforms were generated by synthetic DNA synthesis and restriction enzyme cloning using a mouse KLC1A clone as a starting point. KLC1A coding sequence corresponding to NM_008450.2 was in vector CB6-HA (amino terminal hemaglutinin epitope tag, mammalian expression) flanked EcoRI and BamHI restriction enzyme sites. A unique NheI restriction endonuclease site was added to the KLC1A sequence by site-directed mutagenesis at 1162-1167 by introducing a silent GCCTCC to GCTAGC mutation. This encodes an Ala-Ser sequence within the TPR domain. As the KLC1 isoforms only differ at the carboxy-terminus of the protein, this allowed the synthesis of a series of DNAs (Genscript) that could then be inserted between the newly created NheI site and the existing C-terminal EcoRI site. These sequences were identical to: NM_001025358.2 → NP_001020529.2 for KLC1B; NM_001025359.2 → NP_001020530.2 for KLC1C; NM_001025360.2 → NP_001020531.2 for KLC1D; NM_001025361.2 → NP_001020532.2 for KLC1E; with the exception of KLC1D/E where an unwanted NheI site in the sequence encoding the CTD was removed prior to synthesis by introducing a silent mutation. For expression of the CTDs in bacteria, the DNAs encoding the CTD sequence commencing at Q496 (of QGLD sequence) were amplified by PCR and subcloned into the NdeI/XhoI sites of pET28a. The KLC2 sequence used in this study corresponded to NM_008451.3 → NP_032477.2 (annotated KLC2 isoform 1 in the NCBI Database) and was cloned into the CB6-HA vector using NotI and EcoRI restriction endonuclease sites. The CTD sequence was amplified from Q481 of QGLD sequence and subcloned into pET28a as for KLC1 CTDs. The I560P helix disrupting KLC1E mutant was constructed by PCR-based site-directed mutagenesis using the following primers: forward 5’-CAAACTCCGGGCTTCCCCTAGACGCAGCAGTGAG-3’, reverse 5’-CTCACTGCTGCGTCTAGGGGAAGCCCGGAGTTTG-3’. The L550P helix disrupting KLC2 mutant was constructed by PCR-based site-directed mutagenesis using the following primers: forward 5’-GCTCCGGGATGCTCCGAGACGCAGCAGTGAG-3’, reverse 5’-CTCACTGCTGCGTCTCGGAGCATCCCGGAGC-3’. GFP-tagged KLC1E^CTD^ was cloned from KLC1E^CTD^ cDNA into a CMV driven expression vector (CB6-GFP) for expression in mammalian cells. For expression of GFP-CC-tet-KLC1E^CTD^, cDNA encompassing a *de novo* designed tetrameric coiled coil^39^ and KLC1E^CTD^ was synthesized by Eurofins Genomics (pEX-A128 vector) and subcloned into CB6-GFP. The coiled-coil amino acid sequence was as follows: GELAAIKQELAAIKKELAAIKWELAAIKQ. All plasmids were verified by DNA sequencing.

### Bacterial protein expression and purification

His-tagged KLC^CTDs^ were expressed in Escherichia coli BL21(DE3) cells. Single colonies were picked and grown at 37°C overnight. Small-scale overnight bacterial cultures were used to inoculate two 1-L cultures that were incubated at 37°C until they reached an optical density at 600 nm (OD600) of 0.6 – 0.8. The temperature was then lowered to 16°C and protein synthesis was induced by the addition of 300 μM isopropyl b-D-1-thiogalactopyranoside for 16 h. Cells were harvested by centrifugation at 4,000 g for 30 min at 4°C and resuspended in 25 mM 4-(2-hydroxyethyl)1-piperazineethanesulfonic acid (HEPES) buffer pH 7.4, 500 mM NaCl, 20 mM imidazole, 5 mM β-mercaptoethanol supplemented with protease inhibitor cocktail (Roche). Cell lysis was accomplished by sonication. Insoluble material was sedimented by centrifugation at 13,500 g for 30 min at 4°C prior to loading on His-trap FF columns (GE Healthcare) pre-equilibrated with lysis buffer. Proteins were eluted with an imidazole gradient and fractions containing the target protein collected using an AKTA Prime system (GE Life Sciences). Protein quality was analysed by Coomassie gel analysis. Protein concentration was typically determined by band densitometry using a BSA standard curve due to the low molar extinction co-efficient of the proteins. Proteins were snap frozen and stored at −80°C.

### Purification of HaloTag protein from mammalian cells

The day prior to transfection, 5 x 15 cm dishes were seeded with 4.5×10^6^ 293T cells. Cells were transfected with 20 μg of KHC and KLC expression plasmid (KIF5C/KLC2-TEV-Halo, HA-KLC1B/KLC1E/KIF5C) using the ProFection Mammalian Transfection System (Promega) as per manufacturer’s instructions. Cells were left to express the proteins for 48 h. Cells were then washed with PBS and gently scrapped in 2.5 ml lysis buffer (50 mM Tris (pH 7.5), 150 mM NaCl, 1 mM DTT, 1% Triton-X-100, 0.1% sodium deoxycholate, 1 mM MgCl2, 0.1 mM ADP), supplemented with protease inhibitor cocktail (Promega). Cells were placed on a rocker for 15 min at room temperature before being diluted in 3 volumes of Halo buffer (50 mM HEPES (pH 7.5), 150 mM NaCl, 1 mM DTT, 0.005% IGEPAL CA-630, 1 mM MgCl2, 0.1 mM ADP). The lysate was sonicated, 2 min, 30 sec, pulse for 5 sec on, 10 sec off at 40% amplitude (Vibra Sonics). Insoluble material was pelleted by centrifugation at 10,000 g for 30 min at 4°C. The soluble fraction was incubated with 2.5 ml HaloLink resin slurry (Promega, pre-equilibrated in Halo buffer) and allowed to incubate on a roller for 90 min at room temperature. Resin was washed in batch, by three repeated rounds of centrifugation (1,500 g, 5 min, 4°C) and resuspension in Halo buffer. After the final wash, as much liquid was removed as possible and HaloTEV (Tobacco Etch Virus) protease (300 units, Promega) was added to liberate protein. TEV cleavage proceeded for 60 min at room temperature. The soluble material was separated from the resin by use of a spin filter (ThermoFisher) and centrifugation (5,000 g, 5 min, 4°C). Insoluble material was removed by centrifugation (20,000 g, 5 min, 4°C) and the sample was further purified by size exclusion chromatography on a Superdex200Increase (10/300) column (GE) pre-equilibrated in 20 mM HEPES (pH 7.5), 150 mM NaCl, 1 mM MgCl_2_, 0.1 mM ADP, 5 mM BME. The resulting purified protein was snap-frozen in liquid nitrogen and stored at −80°C until use.

### Circular dichroism

CD spectra were recorded using a Jasco J715 spectropolarimeter (Jasco, MD, USA) with a thermostated cell housing and 1-mm path length cell at 20°C. KLC1D/E synthetic peptides were diluted at 57 μM in PBS buffer pH 7.4 in the absence or presence of DPC (0.35%, Anatrace), DDM (0.05%, Anatrace), OG (2%, Sigma-Aldrich) or LDAO (0.05%, Anatrace). Far UV CD spectra were recorded between 190 and 260 nm. An average of three runs were recorded for each peptide sample.

### NMR spectroscopy

One-dimensional proton spectra of free KLC1D/E synthetic peptide titrated with increasing concentrations of DPC were acquired at ~ 300 mM in PBS buffer supplemented with 10% D2O at 298 K. The DPC-bound peptide sample was prepared by dissolving lyophilized peptide in PBS supplemented with 7% D2O with 0.7% perdeuterated d38-DPC (Cambridge Isotope Laboratories) at a final concentration of ~ 2 mM. Standard two-dimensional NMR spectra (NOESY, TOCSY, COSY, 1H-15N HSQC and 1H-13C HSQC) were acquired at 298 K. NOESY spectra had mixing times of 80, 120 and 250 ms. To guide assignment of the bound peptide, standard two-dimensional NMR spectra (NOESY, TOCSY, COSY, 1H-15N HSQC and 1H-13C HSQC) were also acquired for the free peptide at 278 K. The signals of the free peptide at the lower temperature were better dispersed and sharper than at 298K, allowing partially assignment (Supplementary Figure 5C). All data were collected on a Bruker Avance III HD 700-MHz spectrometer (Bruker, MA, USA) equipped with a 1.7 mm inverse triple resonance micro-cryoprobe using standard pulse sequences from the Bruker library. NMR data were processed in NMRPipe^46^ or Topspin 3.6 and analysed using CCPNMR Analysis v 2.4.2^47^. DSS was used as a chemical shift standard.

### Liposome preparation and co-sedimentation assays

The appropriate lipids were mixed in organic solution and the solvent was evaporated to dryness under a N2 stream. The sample was then kept under N2 stream for an additional 30 min to remove solvent traces. The lipids were swollen in 25 mM HEPES pH 7.4, 150 mM NaCl, 1 mM DTT (dithiothreitol, Sigma-Aldrich) in order to obtain multilamellar vesicles (MLVs) at 1 mg/ml. Large unilamellar vesicles (LUVs) were prepared from MLVs. They were subjected to 10 freeze/thaw cycles, then extruded using 0.2 μm pore size Nuclepore filters (Whatman). Small unilamellar vesicles (SUVs) were obtained by sonicating MLVs with a probe tip sonicator (MSE Soniprep 150, MSE, UK) for 10 min (10 sec on, 10 sec off) on ice. For co-sedimentation assays, 43 μg (unless otherwise specified) liposomes were incubated with 5 μM or 0.2 μM of the indicated KLC^CTD^ or tetrameric kinesin-1, respectively (100 μl final volume) for 30 min and spun down at 170,000 g in a Beckman TLA-100 rotor. Supernatant and pellet fractions were collected and analysed by SDS–PAGE and Coomassie stain.

### PIP arrays

PIP arrays (Echelon Biosciences) were used according to the manufacturer’s instructions. Briefly, strips were blocked with 3% fatty acid-free BSA (Sigma-Aldrich) in PBS buffer containing 0.1% Tween 20 (PBS-T) overnight at 4°C prior to incubation with 1 μg/ml of the indicated protein for 1 h at room temperature. PIP strips were rinsed thoroughly, and protein binding was visualized using an HRP-conjugated anti-His antibody and ECL Chemiluminiscence reagents (Geneflow). Spot intensities were detected on films (GE Healthcare).

### BS3 crosslinking assays

Bis(sulfosuccinimidyl)suberate (BS3) crosslinker (2 mg, Thermo Scientific) was dissolved in Milli-Q water immediately before use and 0-2 mM was added to the corresponding sample. The reaction mixture was incubated at room temperature for 30 min before quenching with sodium dodecyl sulphate (SDS) sample buffer for an additional 15 min. Samples were heated at 95°C for 10 min followed by SDS-PAGE analysis.

### Cryo-electron microscopy

43 μg liposomes were incubated in the presence or absence of 5 μM His-KLC1E^CTD^ for 30 min. Lacey carbon film grids (EM Resolutions) were glow discharged for 30 sec in a Leica ACE 600 Coater system (Leica Microsystems, Heidelberg, Germany). 5 ul of sample were pipetted onto the grids and the sample plunge frozen in liquid ethane using a Leica GP Plunge freezing device (Leica Microsystems, Heidelberg, Germany). The grids were then transferred under liquid nitrogen into a Gatan 626 Cryotransfer system (Gatan, CA, USA) and imaged at ≤ −170°C in a FEI Tecnai 20 LaB6 Scanning Transmission Electron Microscope (FEI, OR, USA) operating at 200kV and using Low Dose imaging conditions. Images were acquired on a FEI CETA 16M CMOS camera (FEI, OR, USA).

### Cell culture, transfection and immunoblotting

HeLa and 293T cells were grown in high-glucose DMEM medium (Sigma-Aldrich) supplemented with 10% FBS (Sigma-Aldrich) and 1% penicillin/streptomycin (Gibco) at 37°C with 5% CO_2_. Transient transfections were performed using Effectene Transfection Reagent (Qiagen) according the manufacturer’s instructions. The efficacy of cell transfection was checked using fluorescence microscopy or western blotting. For immunoblotting, cells grown on 6-well plates were initially washed with ice-cold PBS, then lysed with 100 μl/well of ice-cold radioimmunoprecipitation assay (RIPA) buffer consisting of 50 mM HEPES (pH7.5), 150 mM NaCl, 0.1% NP40, 0.5% sodium deoxycholate and 0.1% SDS supplemented with protease inhibitor cocktail. The homogenates were incubated on ice for 15 min, then cleared by centrifugation at 12,000 g for 15 min at 4°C. Supernatants were collected as soluble fractions. Protein samples were resolved on NuPAGE 4–12% precast gels (Invitrogen), transferred onto PVDF membranes, blocked in 5% milk in TBS-T (20 mM Tris, 0.25 M NaCl, 0.1% Tween-20, pH 7.5 with HCl), and probed with the indicated primary and secondary antibodies (listed above) followed by detection using the Odyssey infrared scanning system (LI-COR Biosciences).

### Immunofluorescence and cell imaging

HeLa cells were seeded on coverslips in 6-well plates prior to transfection. 24 h after transfection, cells were fixed with 4% formaldehyde (Thermo Scientific) for 15 min or ice-cold methanol for 5min. Fixed cells were blocked in 1 % BSA and 4% FCS (Sigma-Aldrich) buffer and incubated for 30 min with primary and secondary antibodies (listed above) diluted in blocking buffer. Cells were DAPI (diamidino-2-phenylindole, Thermo Scientific) stained and placed cell side down in FluorSave reagent (Calbiochem). Confocal microscopy was carried out using a Leica SP5-AOBS confocal laser scanning microscope attached to a Leica DM I6000 inverted epifluorescence microscope (Leica Microsystems, Heidelberg, Germany). A 63x 1.4 NA oil immersion objective (Plan Apochromat BL; Leica Biosystems) and the standard SP5 system acquisition software and detector were used. Image analysis was performed using ImageJ. For co-localisation studies, Pearson’s correlation coefficient was calculated from ~ 40 cells per condition using ImageJ.

### Short interfering RNAs (siRNA) transfection

In HeLa cells, siRNA transfection was carried out through reverse transfection protocol with HiPerFect (Qiagen). 6μl of 10μM siRNAs targeting human KLC1 or KLC2 were mixed with 12μl HiPerFect in high-glucose DMEM medium (Sigma-Aldrich) and combined with 1×10^5^ cells per well of a 6-well plate. Samples were collected 72h after the transfection. For rescue experiments, corresponding mouse KLC1/2 cDNAs (near identical proteins) were transiently transfected so that the siRNA would not recognize the targeting sequence.

### Sequence and secondary structure analysis

Sequence alignment of the KLC^CTDs^ was performed using PROMALS3D^48^ or Clustal Omega using the MegAlign Pro application of the DNAStar Lasergene suite as indicted. Some alignments were further manually edited as indicated in the figure legends. Analysis and characterisation of the predicted secondary structure was performed using JPred4^49^ and Heliquest^50^. Helical 3D models were prepared with PyMol (Schrödinger).

### Kinesin light chain phylogeny

Orthologs from ENSEMBL (PMID 31691826) for KLC1-4 were used as a starting point, followed by sequence similarity searching with BLAST to identify KLC homologues in a wider range of species. Sequences were aligned using the l-INS-I mode in MAFFT^51^, and IQ-TREE2^52^ was used to infer phylogenies under the best-fitting model, selected by BIC. An iterative process of tree inference was used to classify homologous sequences as orthologues and paralogues, and to identify species-specific duplications. For the final tree, the best-fit model was JTT+R6 under the BIC, AIC and AICc criteria.

### Statistical analysis

Data are presented as means ± SEM and were analysed by t-test using Prism 8 (GraphPad Software, CA); *p<0.05, **p<0.01 and ***p<0.001.

---

**Supplementary Figure 1.**
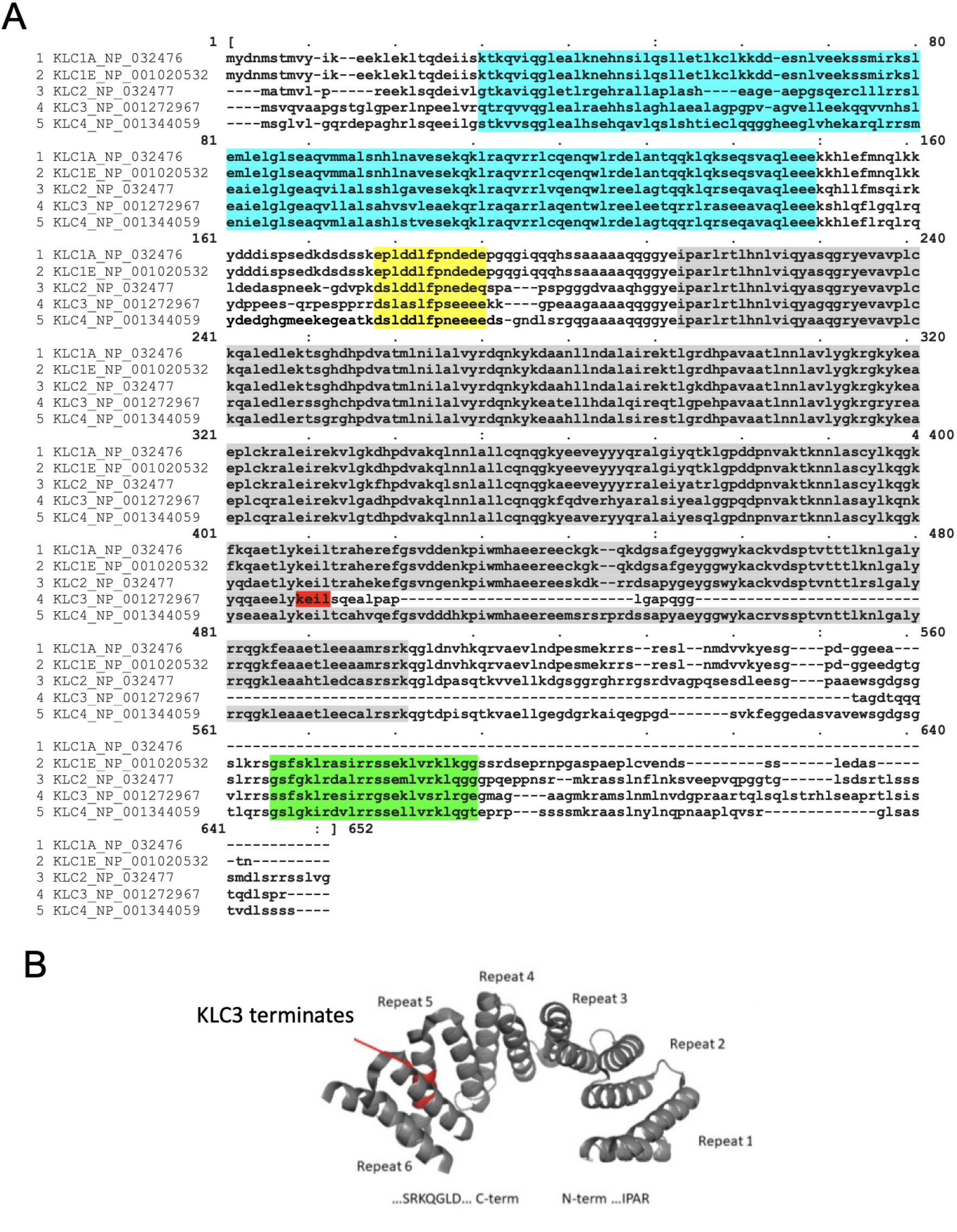
Sequence, domain architecture and structure of the KLCs. (A) Unedited Clustal Omega protein sequence alignment of the indicated KLC paralogues and KLC1 isoforms. Accession numbers are in figure. Predicted and structurally defined features are highlighted in shading and correspond to the domain schematic in Figure 1A. In KLC3, the early truncation of KLC^TPR^ within repeat 5 is highlighted in red and is also shown marked on the structure of KLC1^TPR^ (PDB code 3NF1) in (B).

**Supplementary Figure 2.**
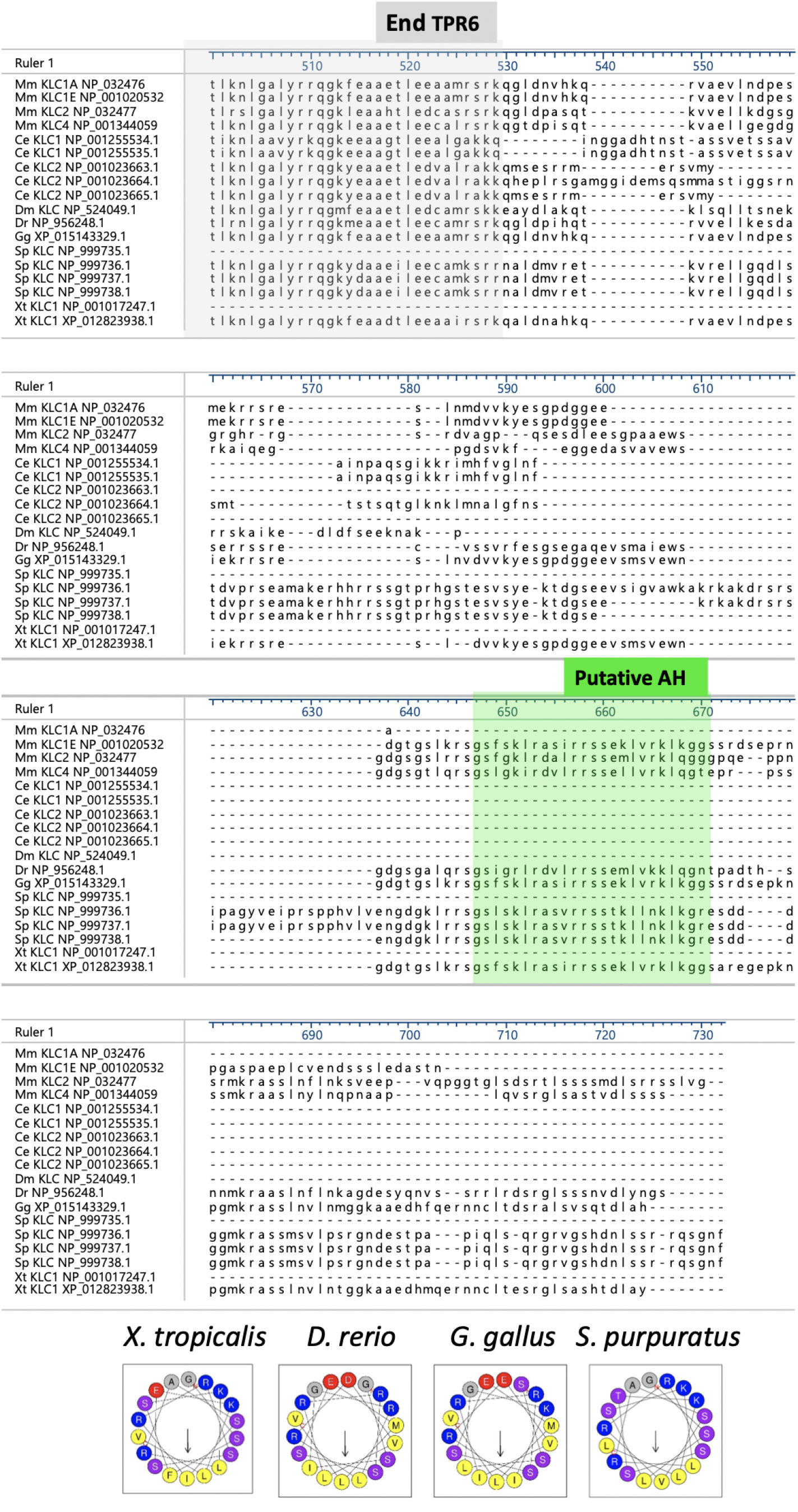
An AH is found in the CTD of deuterostome KLC orthologues. Unedited Clustal Omega alignment of the full length KLC proteins from several species. Figure shows the aligned sequences from the structurally defined end of TPR6 (grey shading) to their carboxy-termini. The well conserved sequence of the AH that is present in the selected deuterostome KLC orthologues, but not in fly or worm, is highlighted in green and also shown as a helical wheel. Mouse KLC1A, KLC1E, KLC2 and KLC4 (as shown in other figures) are compared to selected KLC orthologues identified in the NCBI Protein database. Accession numbers are indicated. Mm – *Mus musculus;* Ce – *Caenorhabditis elegans*; Dm – *Drosophila melanogaster*, Dr – *Danio rerio;* Gg – *Gallus gallus;* Sp – *Stronglyocentrotus purpuratus;* Xt – *Xenopus tropicalis*. Note: as for KLC1 in mammals, in the other species including *Xenopus tropicalis* and *Danio rerio* there also appear to be both shorter and longer forms that are not shown here.

**Supplementary Figure 3.**
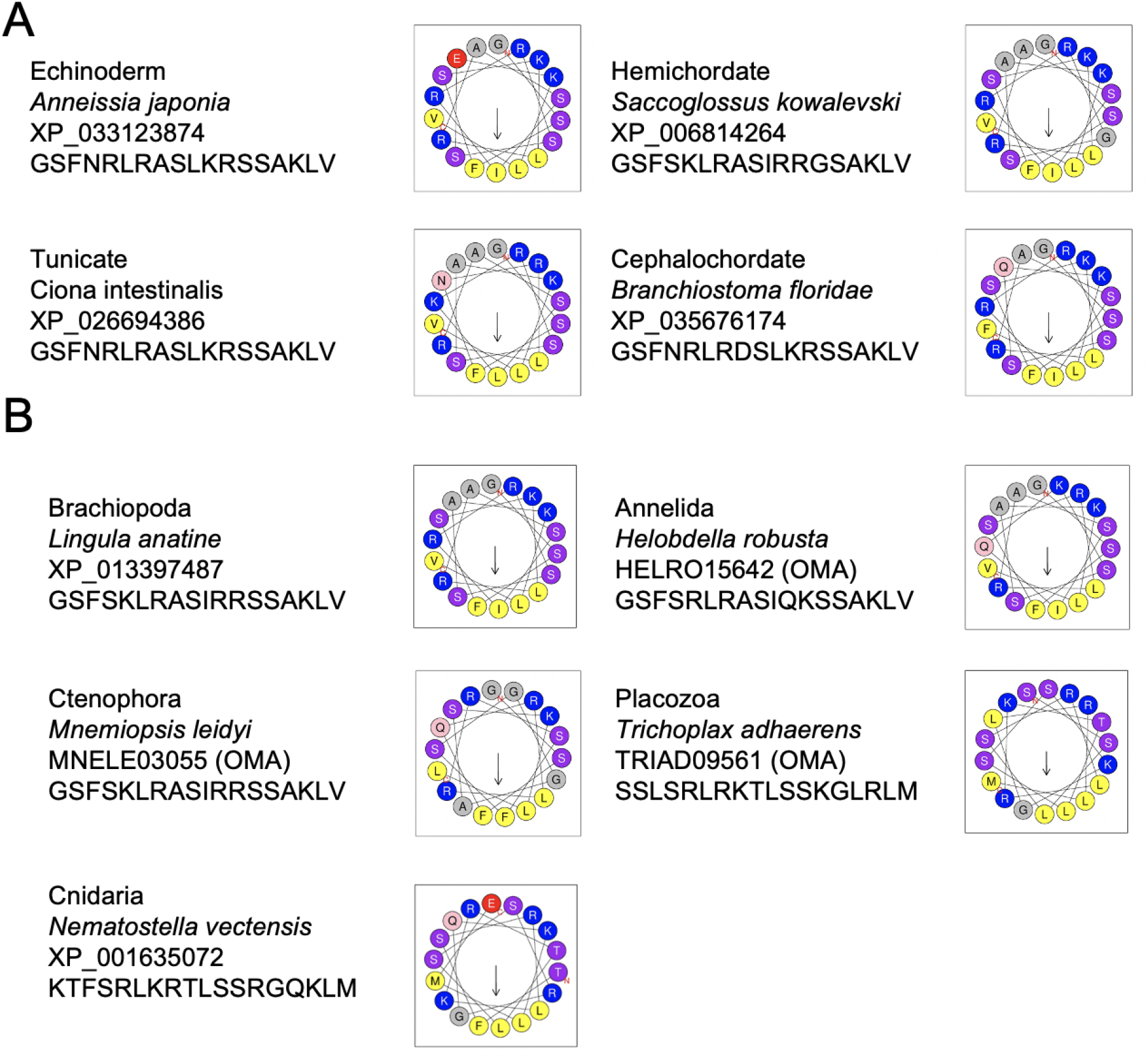
Identification and visualisation of the AH sequence within KLCs of diverse animal species. (A) BLASTp searches were carried out on the non-redundant protein sequences database using the KLC1D/E AH as a query directed by the ‘organism’ constraint, e.g. Echinodermata. Selected deuterostome examples are shown with accession number of the sequence and are represented as helical wheels. (B) Identification of the AH sequence in the CTD of KLC paralogues within a protostome species (*Lingula anatine*) and a non-bilaterian species, the Cnidarian (*Nematostella vectensis*) using KLC^TPR^ as BLASTp query and manual inspection of the subsequent CTD. Further examples in the protostome Annelida (*Helobdella robusta*) and the non-bilaterian Ctenophora (*Mnemiopsis leodyi*) and Placozoa (*Trichoplax adhaerens*) were identified in the OMA database under HOG:0483208 by manual inspection of the CTD sequence combined with Heliquest analysis.

**Supplementary Figure 4.**
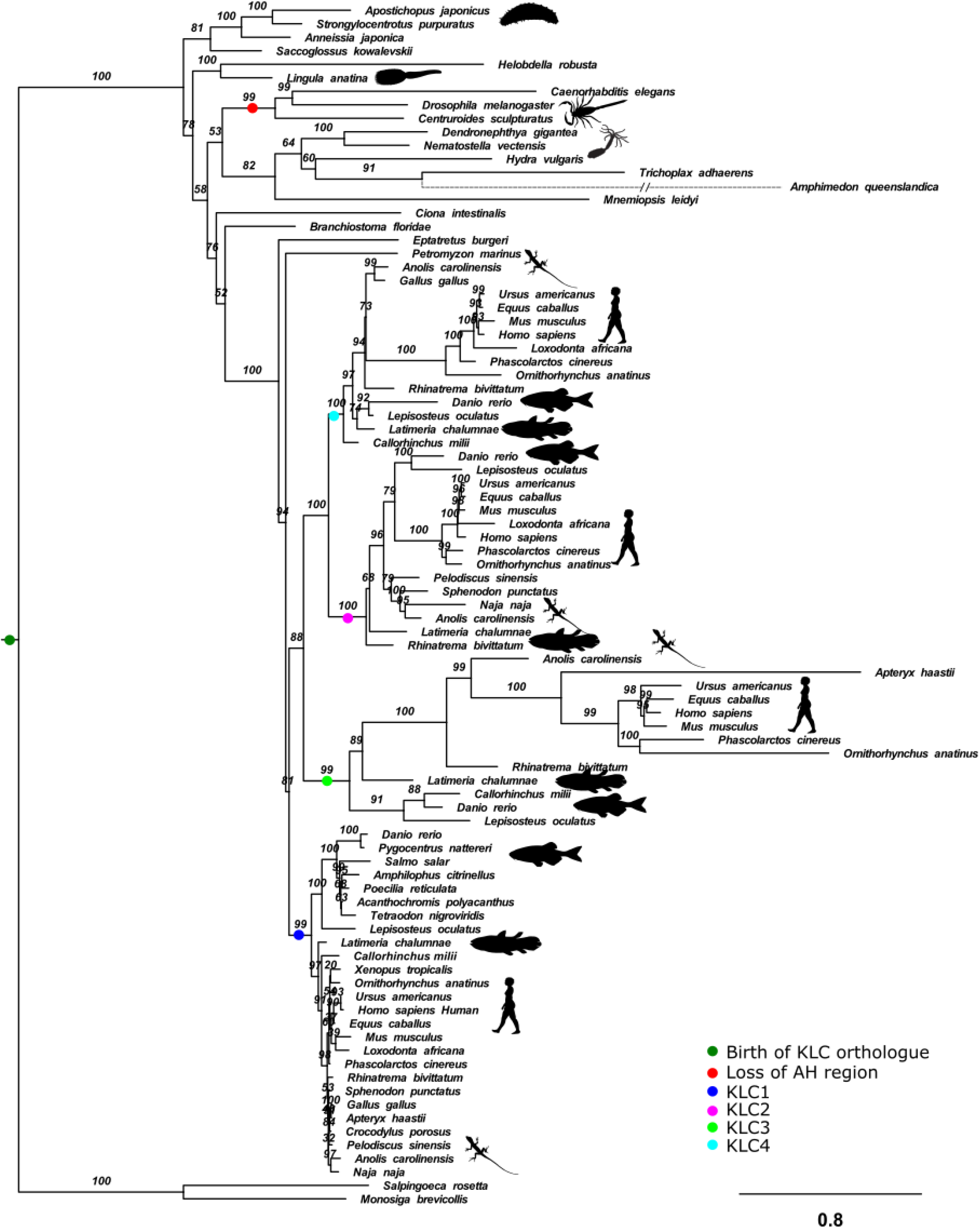
A maximum likelihood phylogeny inferred from orthologous KLC protein-coding genes across Choanozoa. The KLC gene dates back to the common ancestor of animals and choanoflagellates, and the ancestral gene already possessed the amphipathic helix (AH) motif, which was secondarily lost in ecdysozoans. The KLC1-4 paralogues of human and mouse arose from gene duplications in the common ancestor of gnathostomes. The phylogeny was inferred using IQ-TREE2 under the best-fitting substitution model, JTT+R6. Branch supports are ultrafast bootstraps. Branch lengths are proportional to the expected numbers of substitutions per site, as indicated by the scale bar. Copyright: All silhouettes are public domain.

**Supplementary Figure 5.**
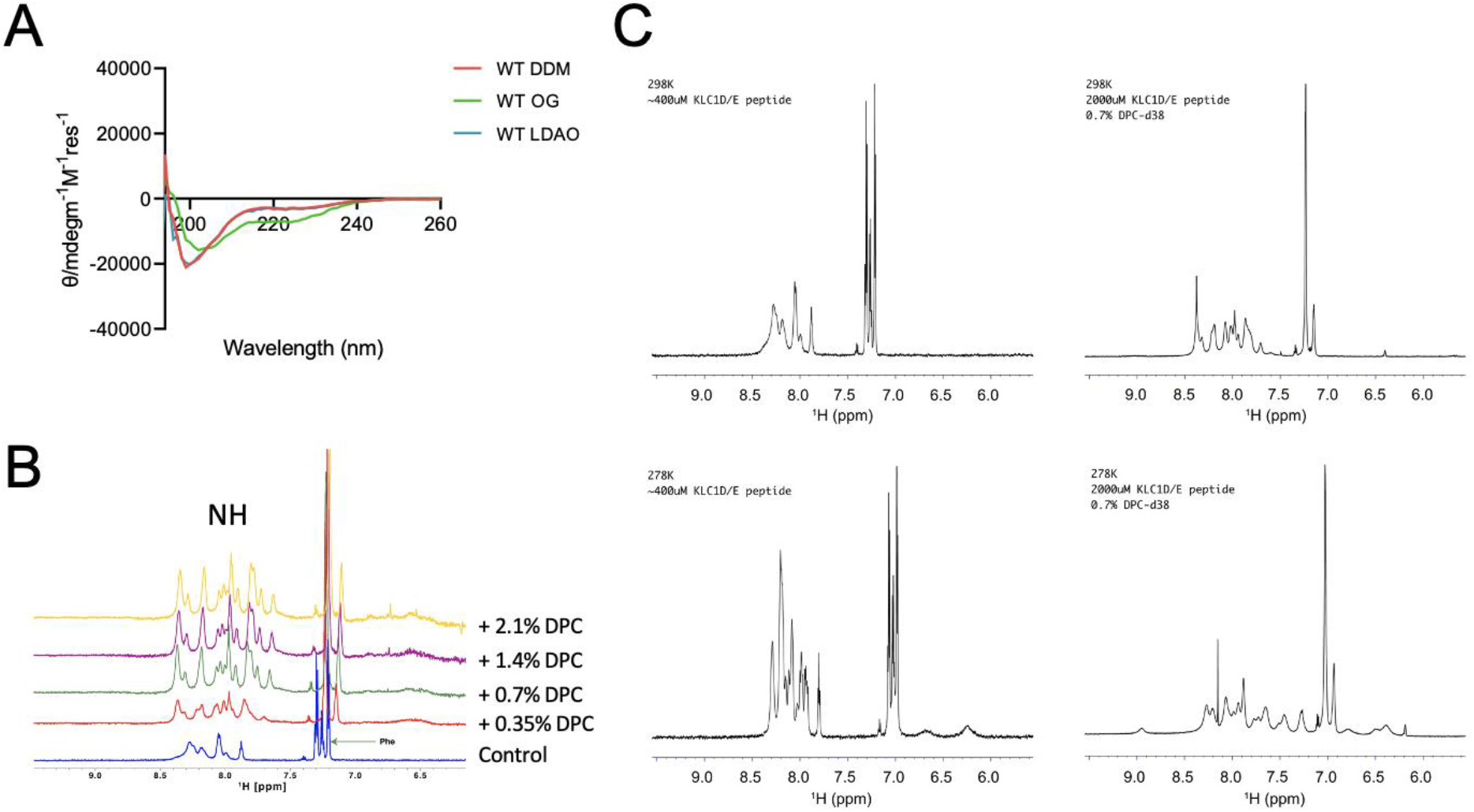
CD and NMR analysis of the KLC1D/E AH peptide. (A) CD spectroscopic analysis of the KLC1D/E synthetic peptide comprising the AH in the presence of 0.05% DDM, 2% OG or 0.05% LDAO. (B) Proton NMR spectroscopic analysis of the KLC1D/E synthetic peptide showing a stacked plot of the NH region in the absence or presence of 0.35, 0.7, 1.4 or 2.1% of DPC at 298K. (C) Proton NMR spectroscopic analysis of the KLC1D/E synthetic peptide at temperatures, peptide concentrations and DPC concentrations indicated on the panels.

**Supplementary Figure 6.**
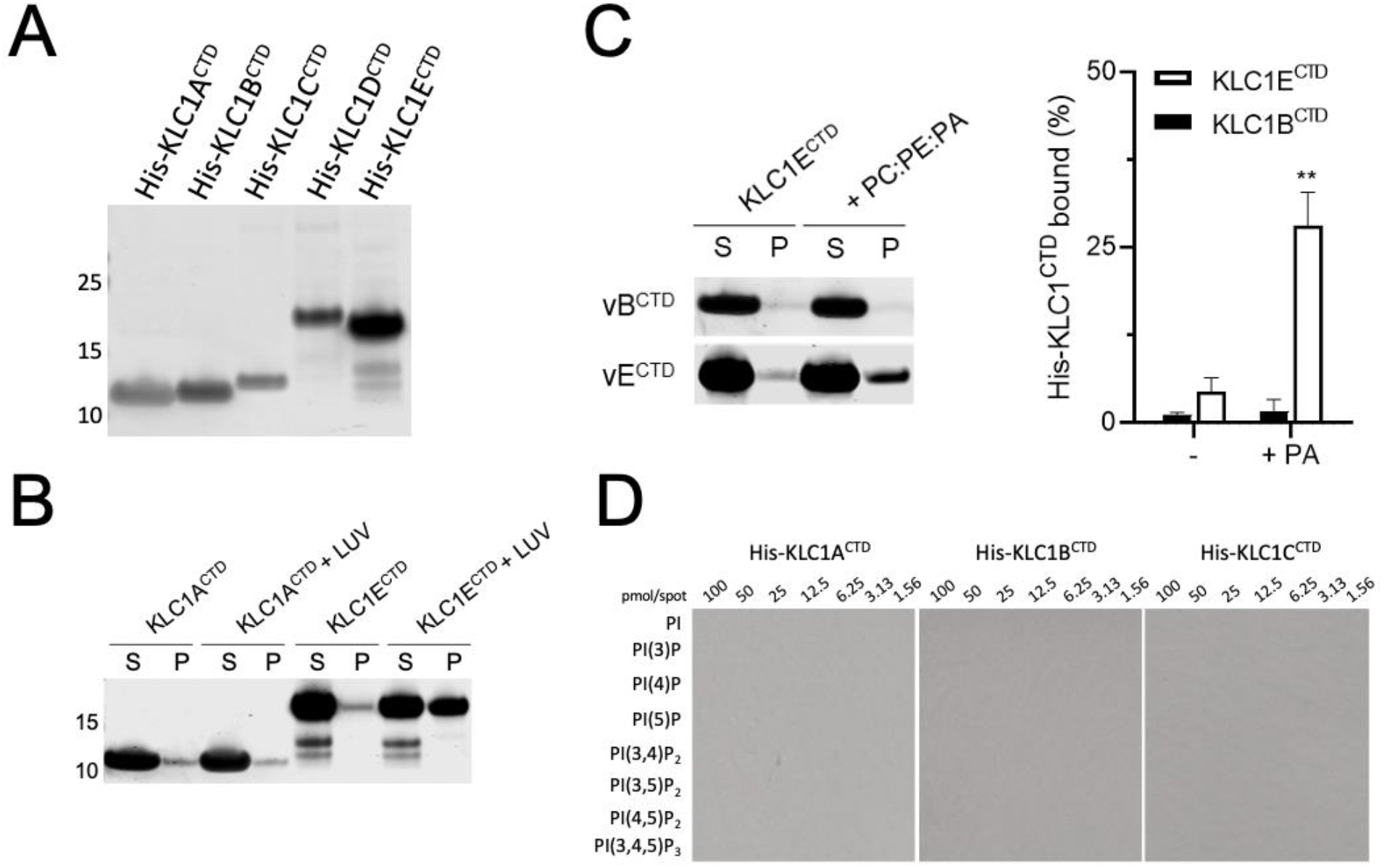
The AH in KLC^CTD^ is required for its direct binding to liposomes. (A) Representative SDS-PAGE and Commassie stain analysis of the recombinant His-tagged KLC1^CTDs^ used in this study. (B) KLC1E^CTD^ but not KLC1A^CTD^ binds membranes in vitro. His-KLC1A/E^CTD^ was incubated with Folch fraction I LUV. The liposome-bound protein fraction was analysed after co-sedimentation by SDS-PAGE stained with Coomassie. (C) His-KLC1B/E^CTD^ was incubated with PC:PE:PA (5:3:2) LUV. The liposome-bound protein fraction was analysed after co-sedimentation by SDS-PAGE stained with Coomassie (left panel) and quantified by densitometry (right panel). Means ± SEM of at least 3 experiments; **p<0.01 compared to the sample without liposomes. (D) His-KLC1A/B/C^CTD^ relative degree of binding to a concentration gradient of eight PI derivatives is shown from a protein lipid overlay assay.

